# Synthetic Genitourinary Image Synthesis via Generative Adversarial Networks: Enhancing AI Diagnostic Precision

**DOI:** 10.1101/2024.05.20.595002

**Authors:** Derek J Van Booven, Cheng-Bang Chen, Sheetal Malpani, Yasamin Mirzabeigi, Maral Mohammadi, Yujie Wang, Himanshu Arora

**Author notes:** **Correspondence:** Himanshu Arora. **Author Contributions:** HA was the chief investigator. DVB, and HA, designed the study and developed the protocol. SM, YM, and MM carried out the pathological analysis plan. HA and DVB coordinated the central modeling investigations. CBC developed Spatial quantification methods. CBC and DVB validated the quantifications. HA, DVB coordinated the data collection. DVB, CBC and HA interpreted the data. HA and DVB developed the first drafts of the manuscript. CBC, SM, MM, YW have accessed and verified all the data in the study. All authors had access to all the data reported in the study. All authors contributed to the review and amendments of the manuscript for important intellectual content and approved this final version for submission. The corresponding author had full access to all the data in the study and had final responsibility for the decision to submit for publication.

## Abstract

In the realm of computational pathology, the scarcity and restricted diversity of genitourinary (GU) tissue datasets pose significant challenges for training robust diagnostic models. This study explores the potential of Generative Adversarial Networks (GANs) to mitigate these limitations by generating high-quality synthetic images of rare or underrepresented GU tissues. We hypothesized that augmenting the training data of computational pathology models with these GAN-generated images, validated through pathologist evaluation and quantitative similarity measures, would significantly enhance model performance in tasks such as tissue classification, segmentation, and disease detection. To test this hypothesis, we employed a GAN model to produce synthetic images of eight different GU tissues. The quality of these images was rigorously assessed using a Relative Inception Score (RIS) of 17.2 ± 0.15 and a Fréchet Inception Distance (FID) that stabilized at 120, metrics that reflect the visual and statistical fidelity of the generated images to real histopathological images. Additionally, the synthetic images received an 80% approval rating from board-certified pathologists, further validating their realism and diagnostic utility. We used an alternative Spatial Heterogeneous Recurrence Quantification Analysis (SHRQA) to assess quality in prostate tissue. This allowed us to make a comparison between original and synthetic data in the context of features, which were further validated by the pathologist’s evaluation. Future work will focus on implementing a deep learning model to evaluate the performance of the augmented datasets in tasks such as tissue classification, segmentation, and disease detection. This will provide a more comprehensive understanding of the utility of GAN-generated synthetic images in enhancing computational pathology workflows. This study not only confirms the feasibility of using GANs for data augmentation in medical image analysis but also highlights the critical role of synthetic data in addressing the challenges of dataset scarcity and imbalance. Future work will focus on refining the generative models to produce even more diverse and complex tissue representations, potentially transforming the landscape of medical diagnostics with AI-driven solutions.

**CONSENT FOR PUBLICATION:** All authors have provided their consent for publication.

## Introduction

Artificial intelligence (AI) has revolutionized the medical imaging landscape, offering innovative applications that aid diagnosis and treatment. In diagnostic radiology, deep learning algorithms, such as those developed by Zebra Medical Vision and Aidoc, analyze X-rays and CT scans to detect a range of conditions, providing faster and sometimes more accurate readings than traditional methods [1-6]. In pathology, companies like PathAI use AI to identify patterns in tissue samples, improving cancer diagnoses [7-11]. Similarly, in ophthalmology, tools like IDx-DR for diabetic retinopathy screening autonomously assess retinal images to identify early signs of disease [4, 12, 13]. In cardiology, AI-powered software like that from Arterys evaluates cardiac MRI and CT scans to provide detailed insights into heart structure and function, aiding in the diagnosis of cardiovascular diseases [14]. Despite these advancements, AI applications are not without concerns. The ‘black box’ nature of many AI systems, where the decision-making process is not transparent, poses challenges in clinical validation and trust. Data privacy and security are also significant issues, as AI models require large datasets for training, potentially exposing sensitive patient information if data is breached or improperly accessed [3, 15, 16]. Real-world breaches, such as the Anthem Inc. and UCLA Health System breaches, underscore these vulnerabilities. Additionally, algorithmic bias and errors in AI systems necessitate meticulous dataset curation and algorithm training to ensure equitable and accurate medical services.

The adoption of Generative Adversarial Networks (GANs) to generate synthetic data presents a promising solution to these challenges [17-19]. GANs can create realistic medical images, reducing the need to use and potentially expose sensitive patient data [17-19]. This method of data augmentation enriches the dataset required for robust AI diagnostic tools and serves as a critical buffer in maintaining patient privacy. In the current study, we utilized GANs for synthetic image generation in genitourinary pathology, highlighting their potential in this context. The GANs underwent rigorous quality control processes, including validation by board-certified pathologists and quantification of image fidelity through Relative Inception Scores and Fréchet Inception Distance, demonstrating high-quality synthetic image production. These images were indistinguishable from real data in many instances, enabling their use in AI diagnostics without the risk associated with actual patient data. By incorporating synthetic data generation via GANs, the healthcare industry can safeguard sensitive patient information, addressing one of the significant cybersecurity concerns of our time. As we continue to navigate the complexities introduced by AI in healthcare, the role of GANs in cybersecurity becomes increasingly pertinent. They represent a promising path forward, integrating AI into medical practice in a secure, ethical, and conducive manner to patient trust and safety.

## Methods

### Cohorts used

We harnessed eight genitourinary tissue types—Bladdar, cervix, kidney, ovary, prostate, testis, uterus, and vagina—obtained from the Genotype-Tissue Expression (GTEx) database, a comprehensive resource that provides open access to tissue expression data. Additionally, histology images from the cancer genome atlas (TCGA) from 500 individuals representing the adenocarcinoma stage were considered as control. Segmentation was performed using PyHIST, a Python-based histological tool, which processed the images into discrete squares of 64, 128, and 256 pixels. Each segment was curated to contain a minimum of 75% tissue content, a criterion set to minimize regional bias and to preserve the representativeness of the histological features.

### Development and Evaluation of a Conditional Generative Adversarial Network

A preliminary conditional Generative Adversarial Network (cGAN) was designed and implemented to assess the performance accuracy of various GAN architectures. The cGAN was developed utilizing Python 3.7.3 and the Tensorflow Keras 2.7.0 package. The generator component of the cGAN comprises three input layers and a single output layer. In parallel, the discriminator component is configured with analogous input, hidden, and output layers. The cGAN’s total parameter count was 7.5 million for each of the evaluated image patterns.

### Implementation and Adaptation of StyleGAN for Tissue Image Analysis

StyleGAN, a progressive generative adversarial network architecture, engineered using Python 3.9 and the TensorFlow framework, leveraging the conditional GAN architecture was used to guide the image synthesis process. This structure allowed the GAN to generate images conditioned on specific tissue types, facilitating targeted image generation. To automate and streamline the process, we employed a bash script tailored for each tissue type that orchestrated the importation of images, their conversion to an RGB color space, and the compilation of these images into a NumPy array. These arrays were then stored as .npy files, ensuring reproducibility and consistency across GAN runs. During the synthetic image generation phase, the generator component of the GAN introduced random noise variables, which were assessed by the discriminator component. This interplay continued iteratively, with the loss graph monitored meticulously until stabilization was observed—a signal to cease the discriminator’s assessment and crystallize the synthetic image output. On average, the GAN system required 2.5 hours per run, yielding a thousand synthetic images per tissue type. The loss functions—mathematical functions quantifying the error between the generated images and the actual images—were pivotal in guiding the GAN’s training. Monitoring these allowed us to fine-tune the GAN’s parameters, with an observed convergence of loss functions around the 182nd epoch. This convergence was deemed the optimal stopping point, indicative of the GAN’s ability to generate images with minimal discrepancy from the target dataset. The term “loss” here referred specifically to the number of images that were not deemed accurate enough by the GAN, thereby being ‘dismissed’ during the iterative training process.

### StyleGAN

The StyleGAN implementation was obtained from NVIDIA labs Github (https://github.com/NVlabs/stylegan) and was run with Python 3.9. Tensorflow version 1.12.0 and CUDA version 10.2 was used. The training images were imported into a TFRecords dataset object and stored as a .tfrecords file. Initial training was performed on a V100 Tesla GPU and took an average of 2.5 days to complete the first round of training. This trained model was then used as the basis for generating new tissue images. The architecture of the StyleGAN was kept exactly as is from the NVIDIA download (https://arxiv.org/abs/1812.04948). The only parameter that was changed to generate sufficient images was the resolution factor. This resolution factor was set to 256 in order to output the images at a quality that could be inspected manually.

### Quantification model

To characterize technical and structural variations between synthetic and real images, we utilized Spatial Heterogeneous Recurrence Quantification Analysis (SHRQA), a robust technique capable of measuring complex microstructures based on spatial patterns[20-22]. The SHRQA process, as shown in Supplementary Figure 4, involves six key steps. It begins with the 2D-Discrete Wavelet Transform (2D-DWT) using the Haar wavelet to reveal patterns not visible in the original image [21-26]. Then, each image is transformed into an attribute vector via the Space-Filling Curve (SFC), which importantly preserves the spatial proximity between pixels in the image within the vector. This step is crucial for analyzing the image’s geometric recurrence in vector form. A trajectory is formed in state space by projecting this attribute vector, highlighting the image’s geometric structure. Through Quadtree segmentation, the state space is divided into unique subregions to discern spatial transition patterns [27, 28]. An Iterated Function System projection is then applied, converting each attribute vector into a fractal plot that represents recurrence within the fractal topology. Finally, these fractal structures are quantified to illuminate the intricate geometric properties of the image, providing a detailed profile.

### Statistical Calculations

FID was implemented in custom scripts developed in house. The FID model was pre-trained using Inception V3 weights for transfer learning. In house code was centered around the FID model and inserted into the StyleGAN to be run during each iteration. Stats were reported at intervals of 1000 and graphed with in house python scripts. Reported FID figures represent an inverse relationship between the images thus the lower our FID figure the more similar the images.

PCA analysis was performed by first transforming the images into numerical arrays. Images were separated into normal and synthetic batches. Intensity was calculated (using the R package imgpalr and magick) as the average of the color of the entire image while keeping the matrix framework (ie positional arguments were retained). PCA was conducted using the general prcomp function in R and plotted results were displayed in ggplot2.

### Data Sharing

De-identified participant data will be made available when all primary and secondary endpoints have been met. Any requests for trial data and supporting material (data dictionary, protocol, and statistical analysis plan) will be reviewed by the trial management group in the first instance. Only requests that have a methodologically sound proposal and whose proposed use of the data has been approved by the independent trial steering committee will be considered. Proposals should be directed to the corresponding author in the first instance; to gain access, data requestors will need to sign a data access agreement.

## Results

### GAN Model Selection

To evaluate the performance of various GAN architectures and select the most appropriate one, n= digital histology images were downloaded from Genotype-Tissue Expression (GTEx) database for Prostate ----. Image patches extracted from -- individuals were divided into training cohorts. Each training cohort was subjected to cGAN, StyleGAN, and dcGAN architectures [29-31]. A total of --synthetic images generated by each GAN were fed into a generic CNN for classification. The cGAN achieved an accuracy of ---, while the StyleGAN and dcGAN demonstrated accuracies of --- and ---, respectively. Although the StyleGAN and dcGAN exhibited similar accuracies, the quality of output was more extensive for StyleGAN, which is particularly important considering the less amount of heterogeneity that exists in standard/non-cancer tissue image types.

### Image synthesis

Once the GAN was selected, the GTEx database was used to extract digital histology images from eight genitourinary tissue types; 129 images were available for Bladdar, 81 for Cervix, 599 for Kidney, 252 for Ovary, 599 for the Prostate, 588 for Testis, 234 for Uterus and 272 for Vagina. Several factors, such as staining protocols, tissue quality, section thickness, tissue folding, and the amount of tissue on the slide, could negatively impact the efficiency of GAN model in generating high quality data [32]. To account for this, we conducted pre-processing normalization of the images. Specifically, we selected all the images from all tissue types and evaluated their color distribution by calculating the mean value of RGB colors and normalizing it. Images with an RGB mean intensity value two standard deviations away from the total mean value of all samples were identified as outliers and removed from the dataset. In total, --- images were discarded due to being outliers. Overall, our pre-processing steps helped to reduce the variability in tissue biopsy images and ensured a more consistent training dataset for the StyleGAN model. Post-processing, these images were used to train the StyleGAN model. The network generator created a total of 200 random synthetic images for each of the tissue type. The patch size of each of these images was set at 5000*5000 to allow sufficient quality for the pathologist evaluation. These image patches were analyzed using the Adam optimization algorithm. This process helped us find the best iteration value for our model, which was 15,000 iterations. 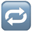Figure 1a summarizes the steps in the processing and generation of synthetic images, and Figure 1b and Supplementary Figures 1-8 showcase the examples of synthetic images generated from eight GU tissues.

**Figure 1.**
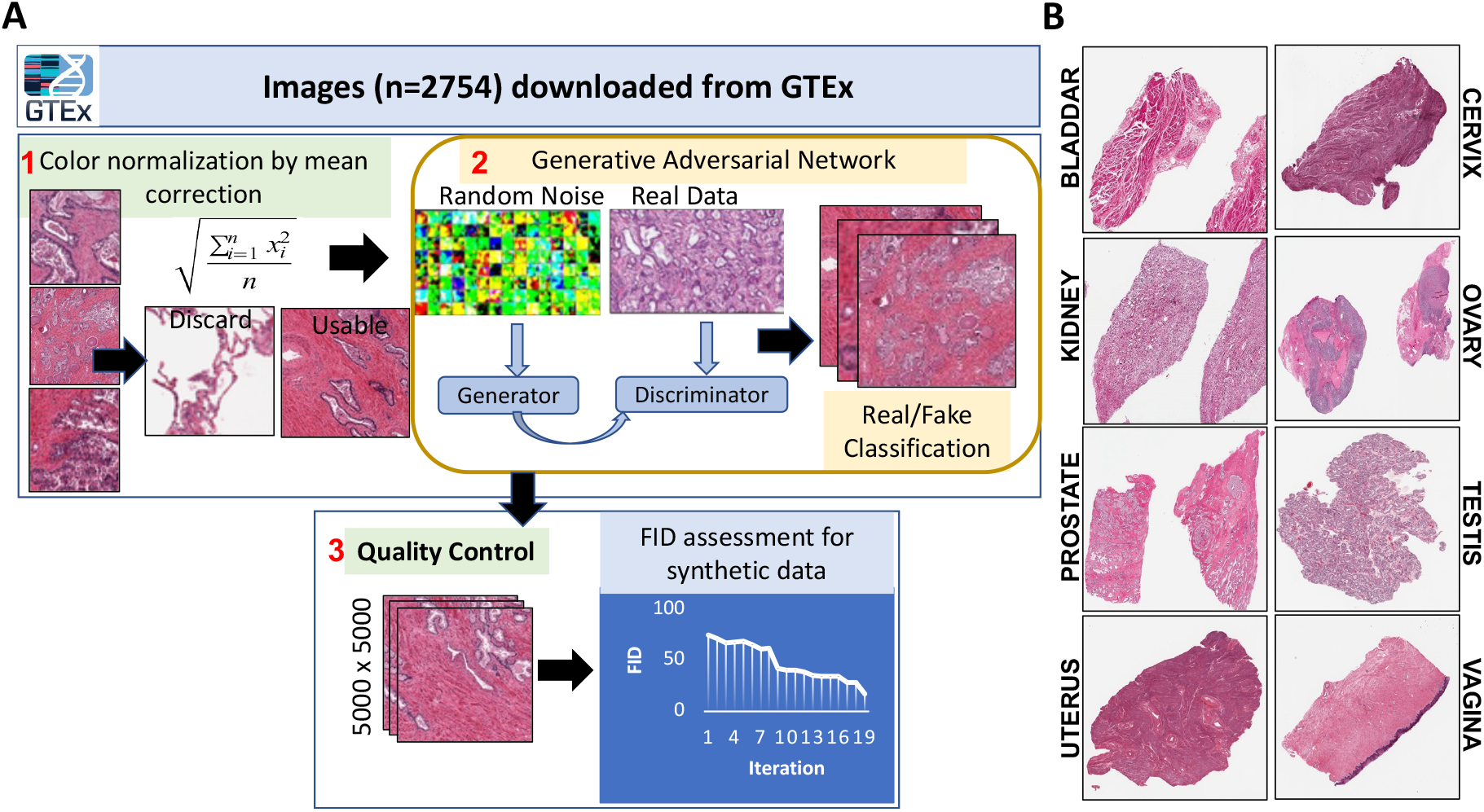
(A) GAN Workflow. Images were normalized, run through the GAN, and then put through QC. (B) Synthetic images were generated for each GU tissue type respectively.

Next, we applied standard machine learning metrics to evaluate the synthetic images. The Relative Inception Score (RIS) was a primary metric, measuring the clarity and variety of the generated images. A high RIS of 17.2 with a remarkably low standard deviation of 0.15 across different tissue types demonstrated the synthetic images’ consistent quality. Furthermore, the Fréchet Inception Distance (FID), a crucial index for GAN performance, was used to compare the distribution of generated images with real images. An FID score that stabilized at 120 indicated that the synthetic images closely mirrored the distribution of the real tissue images, solidifying the efficacy of our GAN model.

### Quality Control Through Expert Evaluation

The synthetic images underwent a rigorous review process for quality control. A subset of synthetic prostate images were subjected to detailed visual inspection focusing on aspects such as sharpness and resolution. This scrutiny was critical to ensure that the generated images met the high standards required for clinical use. For this, two certified pathologists conducted an independent review of the synthetic images cohorts, where they were provided with a randomized pool of 20 images per tissue type, consisting of a mixture of 15 synthetic and five real images, totaling 160 images. The pathologists were tasked to evaluate the quality of the images and highlight the concerns they may have for each tissue type. Table 1 summarizes the quality evaluation outcomes of pathology evaluation for all eight tissue types. Supplementary Figure S1-4 shows the 20 images per tissue type shared with the pathologists. Results highlighted an 80% approval rate, signifying a robust endorsement of the synthetic images’ clinical utility.

**Table 1.**
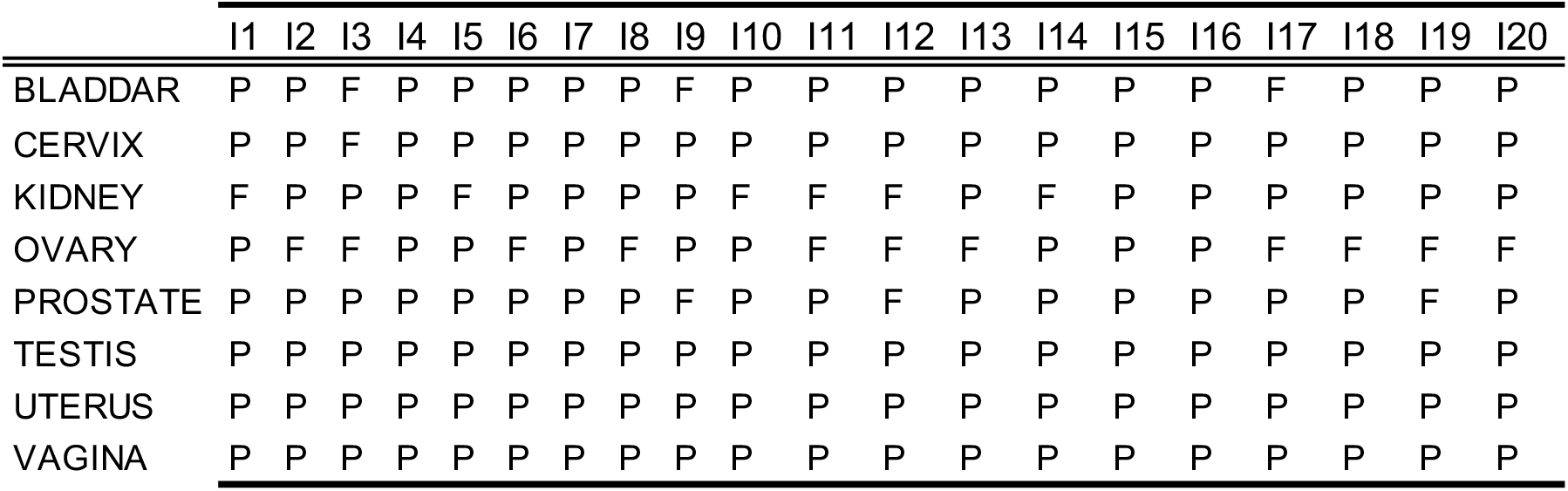
Outcomes of image quality assessment of 20 images per tissue type by two pathologists. The pathologists were subjected to query of “QC Pass” (P) or “QC Fail” (F), and to highlight any concerns they had during the evaluation process.

|  | Image Quality Assessment by Pathologist |  |  |  |  |  |  |  |  |  |  |  |  |  |  |  |  |  |  |  |
| --- | --- | --- | --- | --- | --- | --- | --- | --- | --- | --- | --- | --- | --- | --- | --- | --- | --- | --- | --- | --- |
|  | I1 | I2 | I3 | I4 | I5 | I6 | I7 | I8 | I9 | I10 | I11 | I12 | I13 | I14 | I15 | I16 | I17 | I18 | I19 | I20 |
| BLADDAR | P | P | F | P | P | P | P | P | F | P | P | P | P | P | P | P | F | P | P | P |
| CERVIX | P | P | F | P | P | P | P | P | P | P | P | P | P | P | P | P | P | P | P | P |
| KIDNEY | F | P | P | P | F | P | P | P | P | F | F | F | P | F | P | P | P | P | P | P |
| OVARY | P | F | F | P | P | F | P | F | P | P | F | F | F | P | P | P | F | F | F | F |
| PROSTATE | P | P | P | P | P | P | P | P | F | P | P | F | P | P | P | P | P | P | F | P |
| TESTIS | P | P | P | P | P | P | P | P | P | P | P | P | P | P | P | P | P | P | P | P |
| UTERUS | P | P | P | P | P | P | P | P | P | P | P | P | P | P | P | P | P | P | P | P |
| VAGINA | P | P | P | P | P | P | P | P | P | P | P | P | P | P | P | P | P | P | P | P |

### Geometric Analysis of Image Characteristics

To delve deeper into the geometric properties of the synthetic images, we employed Spatial Heterogeneous Recurrence Quantification Analysis (SHRQA). This involved initial image preprocessing, including grayscale conversion, noise reduction, contrast enhancement, and normalization, to reduce the unrelated noise and amplify underlying patterns within the images. Subsequently, each image was transformed into an attribute vector through the application of a Hilbert Space-Filling Curve, a technique that preserves the spatial proximity relationships of the image pixels in a one-dimension vector. This vectorization facilitated a detailed analysis of the geometric recurrence and structural intricacies within the images. By applying the Iterated Function System projection, we were able to identify and quantify recurrent fractal structures, thereby providing a robust profile of the images’ geometric fidelity (Figure 2).

**Figure 2.**
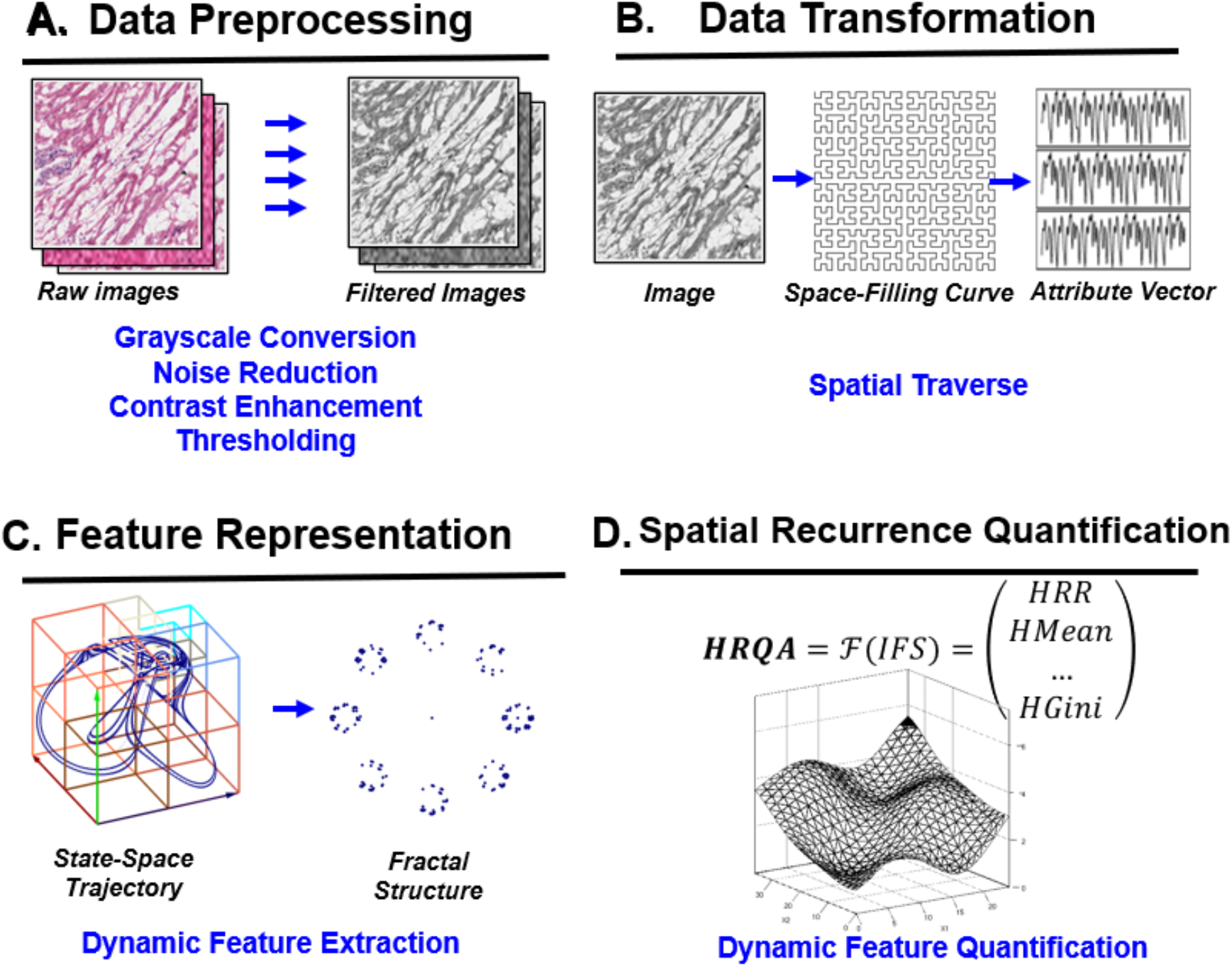
The framework of the Spatial Heterogeneous Recurrence Quantification Analysis (SHRQA). Initially, each image undergoes standard image preprocessing including grayscale conversion, noise reduction, contrast enhancement, and thresholding. This amplifies intricate patterns and minimizes environmental noise. Subsequently, a space-filling curve transforms each image into an attribute vector, preserving the majority of its proximity information. Through state-space construction, pixel color/intensity transitions form a trajectory in the state space. These transitions are then projected into an Iterated Function System (IFS) to capture complex dynamic properties. The image’s nuanced geometric properties are then mathematically described using recurrence quantification analysis. Ultimately, the extracted spatial recurrence characteristics can be employed to profile images.

The SHRQA method was first applied to examine the spatial recurrence properties of real and synthetic image patches across test tissue data which was prostate in this specific scenario. Our sample set included an equal number of patches from real and synthetic sources, with a balanced representation of each phenotype. To add an extra layer of validation, we downloaded histology images from the cancer genome atlas (TCGA) from individuals representing the adenocarcinoma stage. These images, representing different stages of cancer progression (represented by Gleason grade), were randomized.

On all the image types (normal-original (NO), normal-synthetic (NS), and cancer-original (CO)), Segmentation was performed using PyHIST, a Python-based histological tool, which processed the images into discrete squares of 256 pixels. Each segment was curated to contain a minimum of 90% tissue content, a criterion set to minimize regional bias and to preserve the representativeness of the histological features. We analyzed 2000 image patches, each 256×256 pixels, evenly split between real and synthetic. SHRQA quantitatively outlined each patch’s microstructures. From an initial extraction of 112 spatial recurrence features per patch, LASSO selected 102 features as significant to the Gleason pattern. Hotelling’s T-squared test, a multivariate extension of the two-sample t-test, compared the spatial recurrence attributes of real versus synthetic patches. The resulting p-values of 0.4039 signified no significant differences in spatial recurrence properties between NO and NS, but a significant difference was observed between NO and CO (p= 1.353e-07), NS and CO (p= 1.759e-07). as confirmed by the T-squared tests’ p-values for each Gleason pattern.

We also employed PCA on the spatial recurrence properties [33-39], visualized using radar charts, revealing that the top ten principal components capture 90% of the variability. This allowed us to map the distributions of spatial properties for real and synthetic images across phenotypes, as depicted in Figure 3. Notably, while distributions aligned closely between real and synthetic images (NO-NS), significant differences were evident between NO-CO and NS-CO. These findings across image sections validate the model’s efficiency in capturing the geometric intricacies consistent with real images.

**Figure 3.**
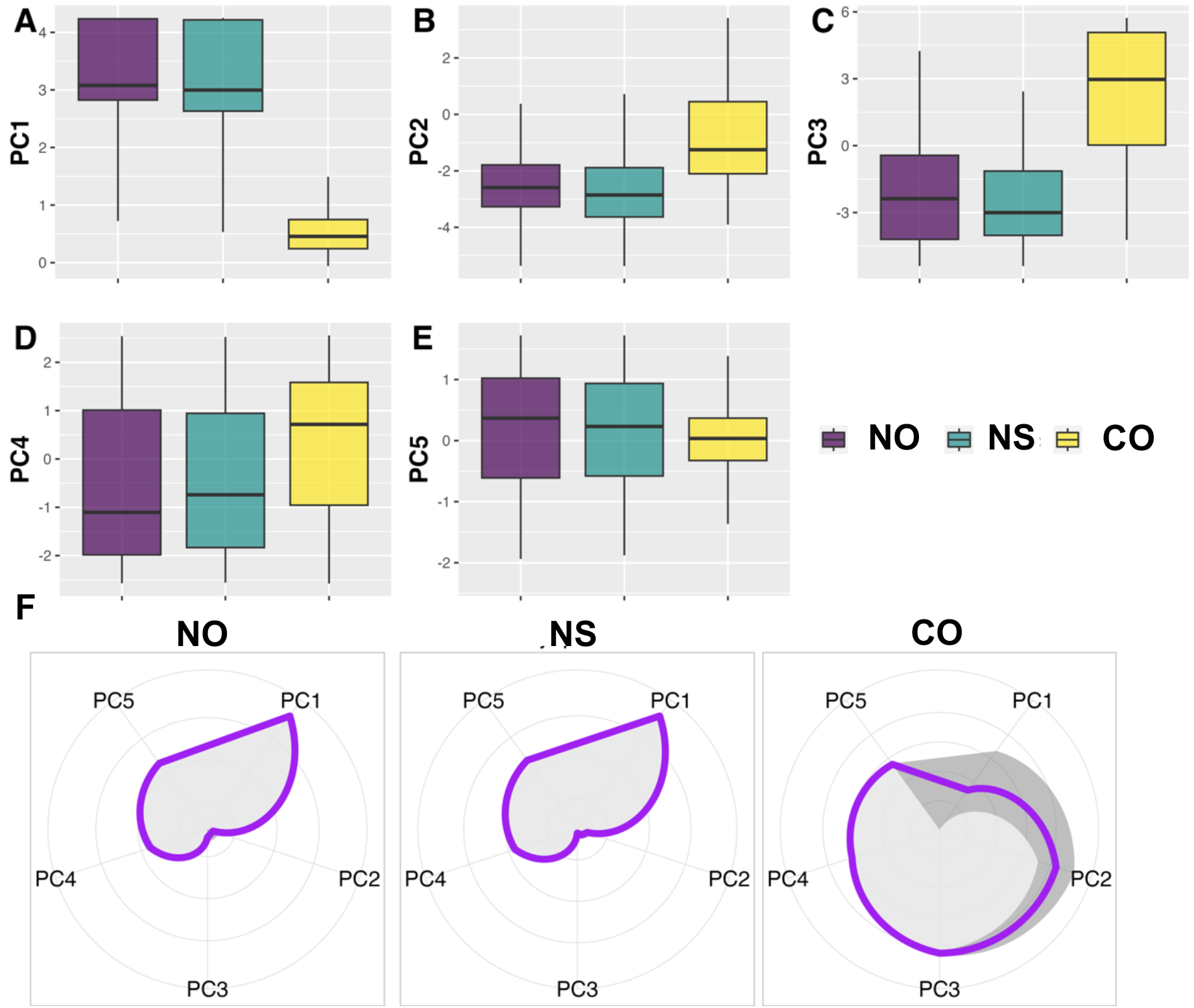
(A) The comparison of spatial recurrence properties between normal-original (NO), normal-synthetic (NS), and cancer-original (CO), on the first four PCs (contain >95% of data variability). The distributions of this four PCs are similar between real and synthetic. (B) The distributions of spatial recurrence properties (in the first 5 Principal Components (PCs), which contain > 95% of data variability) underlying different patterns for both real and synthetic patches. Note that the purple lines indicate the mean values of each feature, and the gray area shows the 95% confidence interval. Our results indicate that while the distributions of spatial properties are closely aligned between NO and NS, they markedly differ when comparing CO.

## Discussion and Conclusion

The application of Generative Adversarial Networks (GANs) in producing synthetic medical images, as demonstrated by our research, has significant implications for healthcare. By generating synthetic images that are virtually indistinguishable from real histological samples, GANs provide a powerful tool for training AI systems without the risk of exposing sensitive patient information. This is a key consideration given the notable cybersecurity incidents in recent years, such as the Anthem Inc. and UCLA Health System breaches, which exposed the data of millions. Our study’s success in generating high-quality synthetic genitourinary images serves as a proof of concept for the broader application of GANs in medical imaging. By employing this technology, healthcare providers can enhance the robustness of AI diagnostic tools while maintaining stringent data security. For instance, rather than relying on vast databases of patient images, which pose a potential risk if compromised, medical AI applications can be trained using synthetic datasets that carry no privacy concerns.

The practicality of synthetic images generated by GANs is further supported by their performance in standard machine learning metrics and approval by expert pathologists. This dual validation underscores the potential of GANs not only in generating training data but also in providing a buffer against data breaches. As AI continues to permeate the medical field, the ability to create diverse, high-fidelity datasets through GANs becomes increasingly valuable, offering a safeguard against the risks associated with the collection and storage of large-scale patient data.

Looking ahead, the expansion of this methodology to other tissue types and medical conditions could revolutionize the field of medical diagnostics. For example, AI models trained on GAN-generated images could support the early detection of rare diseases without requiring access to potentially sensitive real-world data. Similarly, the generation of synthetic images for rare pathologies could aid in developing diagnostic models where real data is scarce or difficult to obtain due to privacy concerns. However, there are limitations that needs to be addressed for appropriate application of the generated data. For example, validation of synthetic data, though it was beyond the scope of the current study, should be done by using various publicly available and developed convolutional neural network (CNN) models. Another limitation is the time the StyleGAN takes to generate the synthetic data can limit its widespread application. A potential solution to this is to generate image patches of smaller sizes, which may introduce a reduction in the quality of the data but would significantly increase the model’s efficiency. Third, the quantification models utilized in this study may benefit from assisted learning modules, which will allow feature-specific quantification with respect to each tissue type, unlike its current stage.

In conclusion, the implementation of GANs in digital pathology represents a promising avenue for enhancing both the effectiveness of AI in medical diagnostics and the security of patient data. As healthcare continues to evolve alongside AI, the development of secure, synthetic datasets through GANs will be crucial in mitigating the risks of data breaches while unlocking the potential for more advanced, personalized treatment options.

---

**Supplementary Figure 1.**
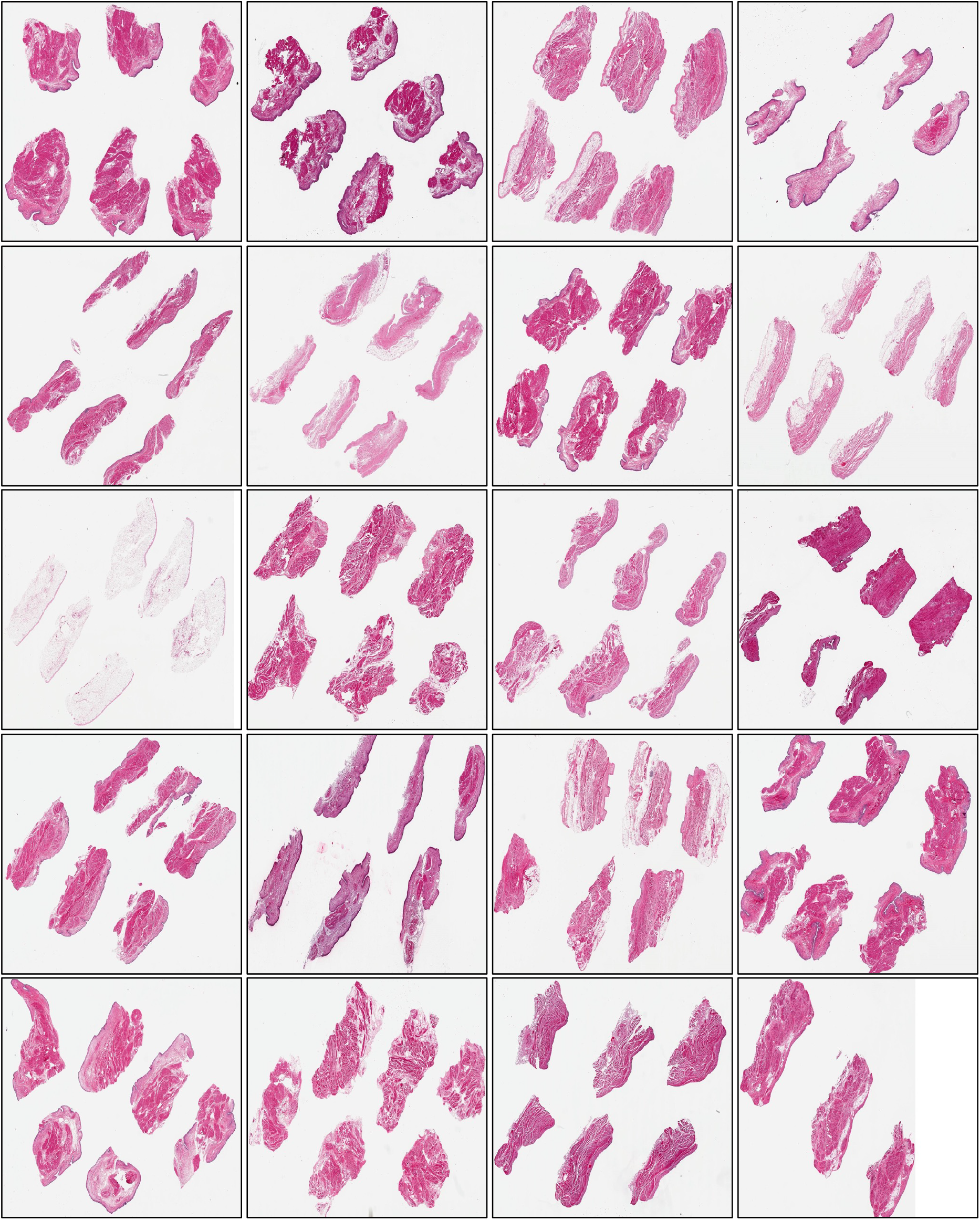
Showing synthetic images of Bladdar.

**Supplementary Figure 2.**
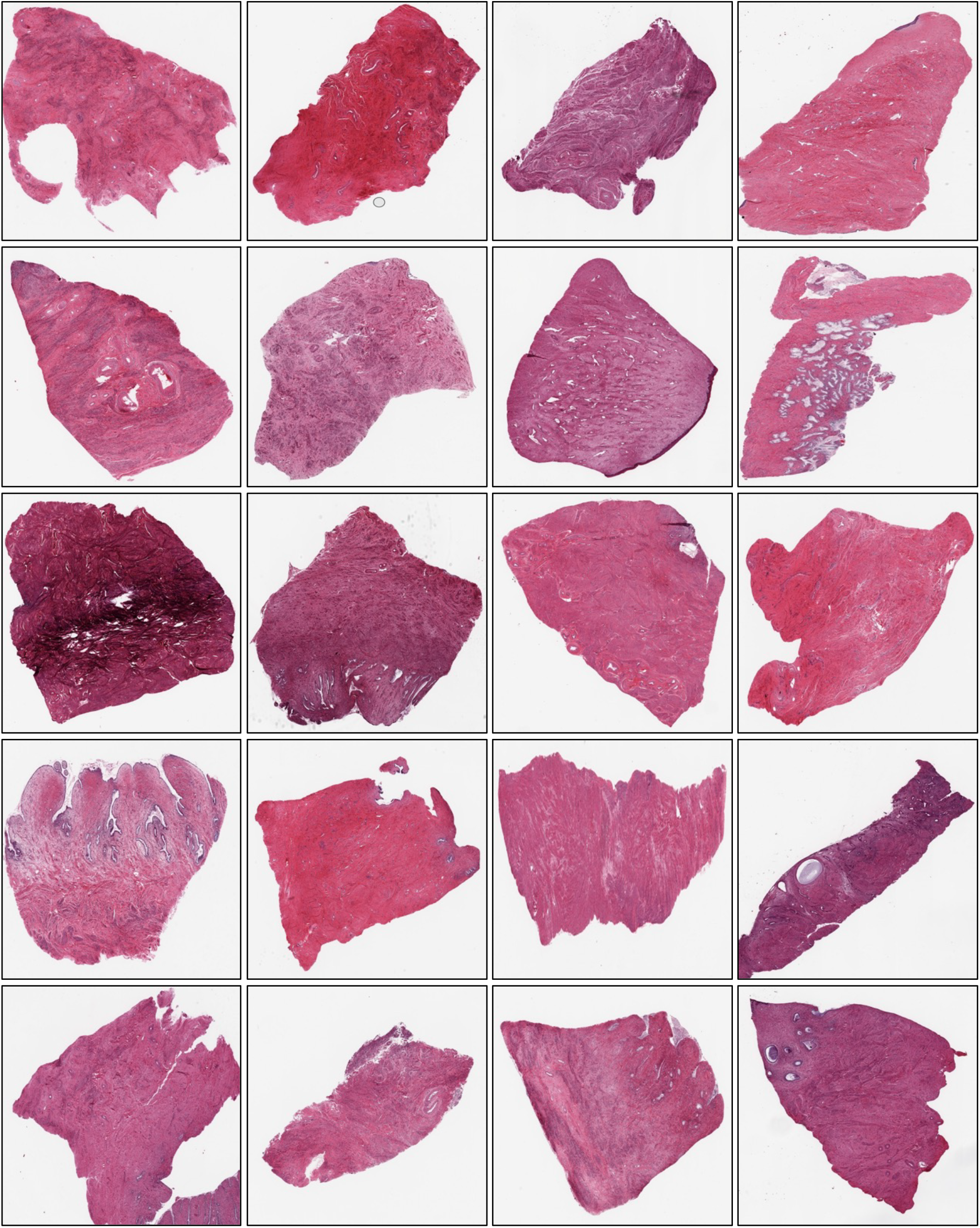
Showing synthetic images of the Cervix

**Supplementary Figure 3.**
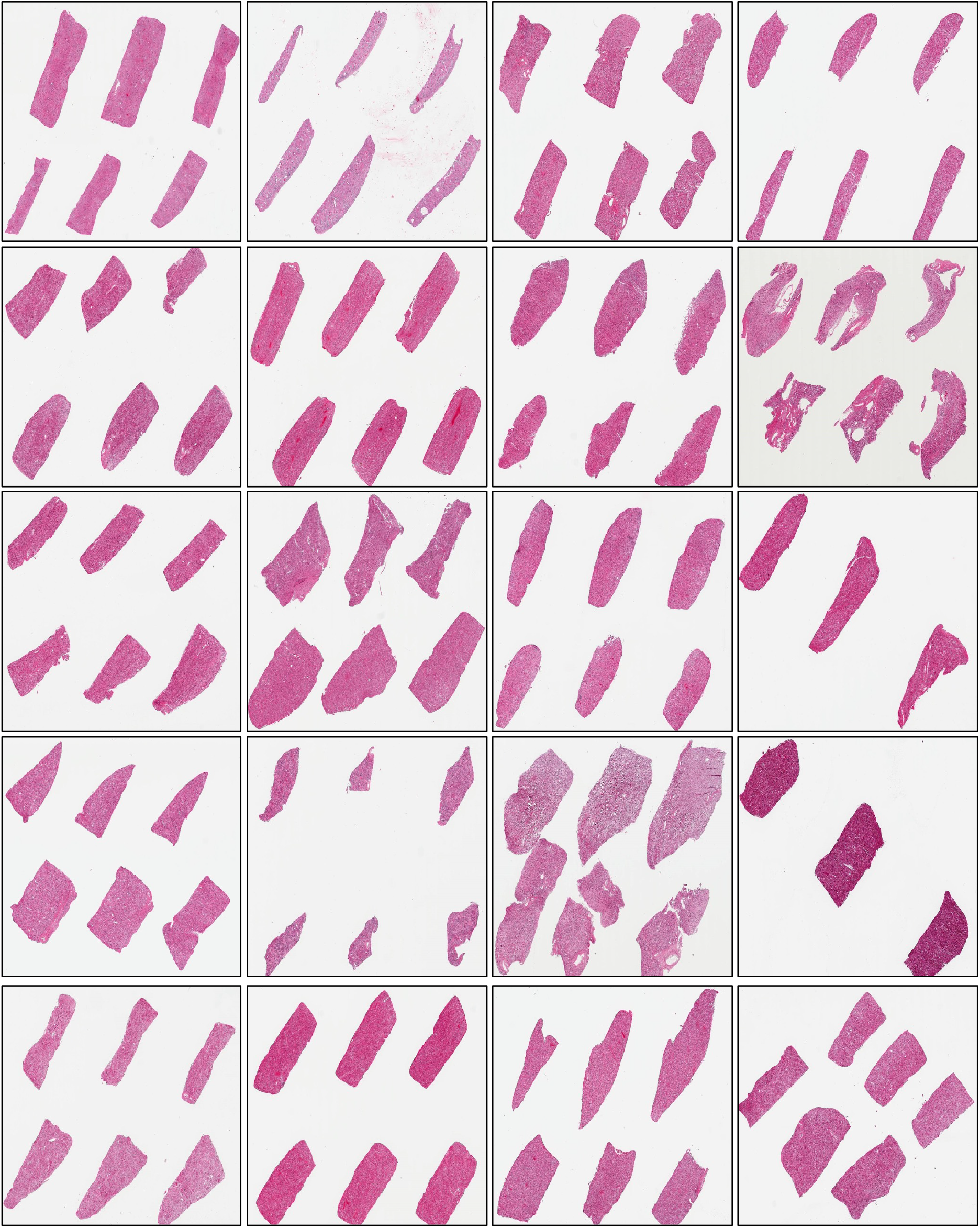
Showing synthetic images of the Kidney.

**Supplementary Figure 4.**
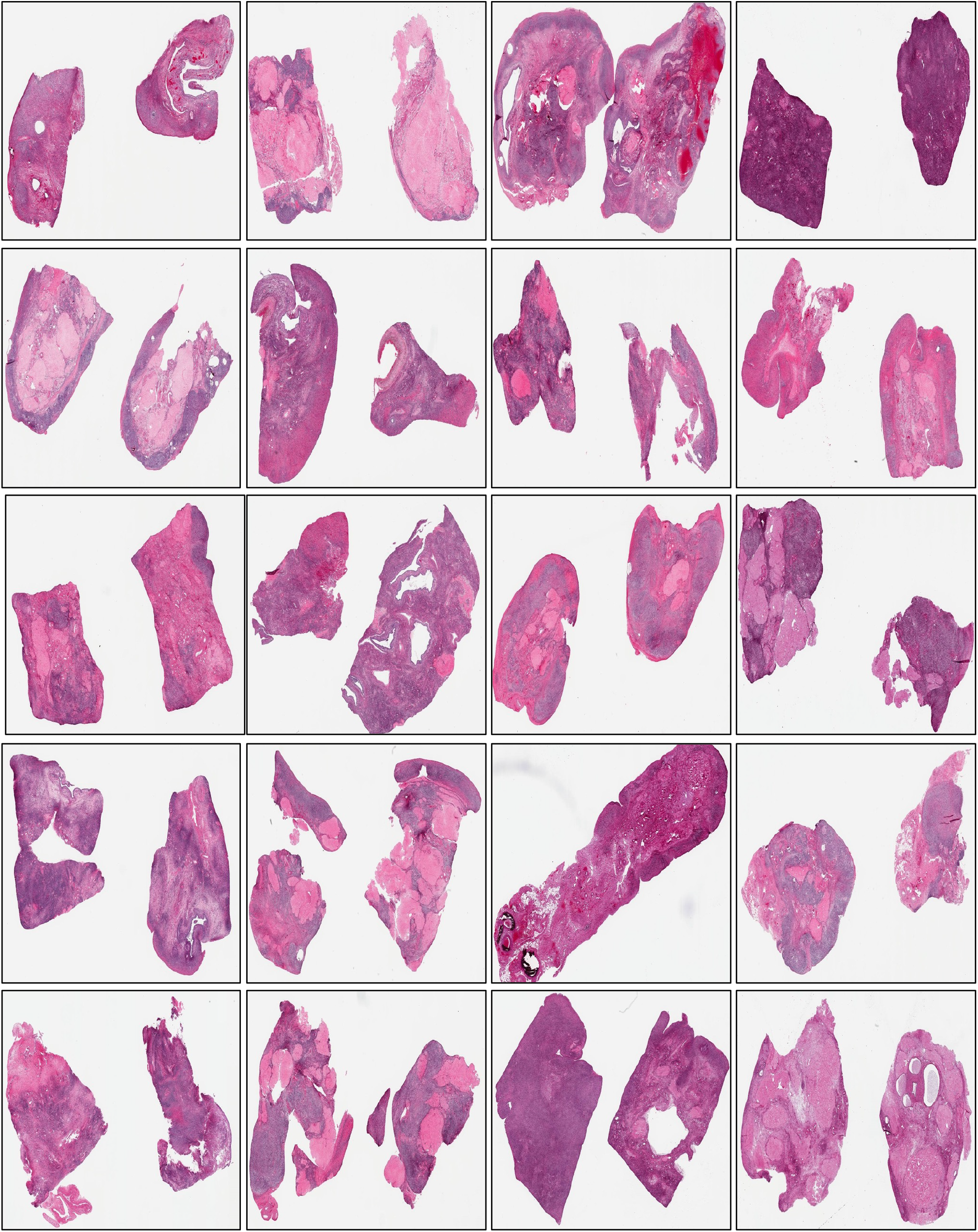
Showing synthetic images of the Ovary.

**Supplementary Figure 5.**
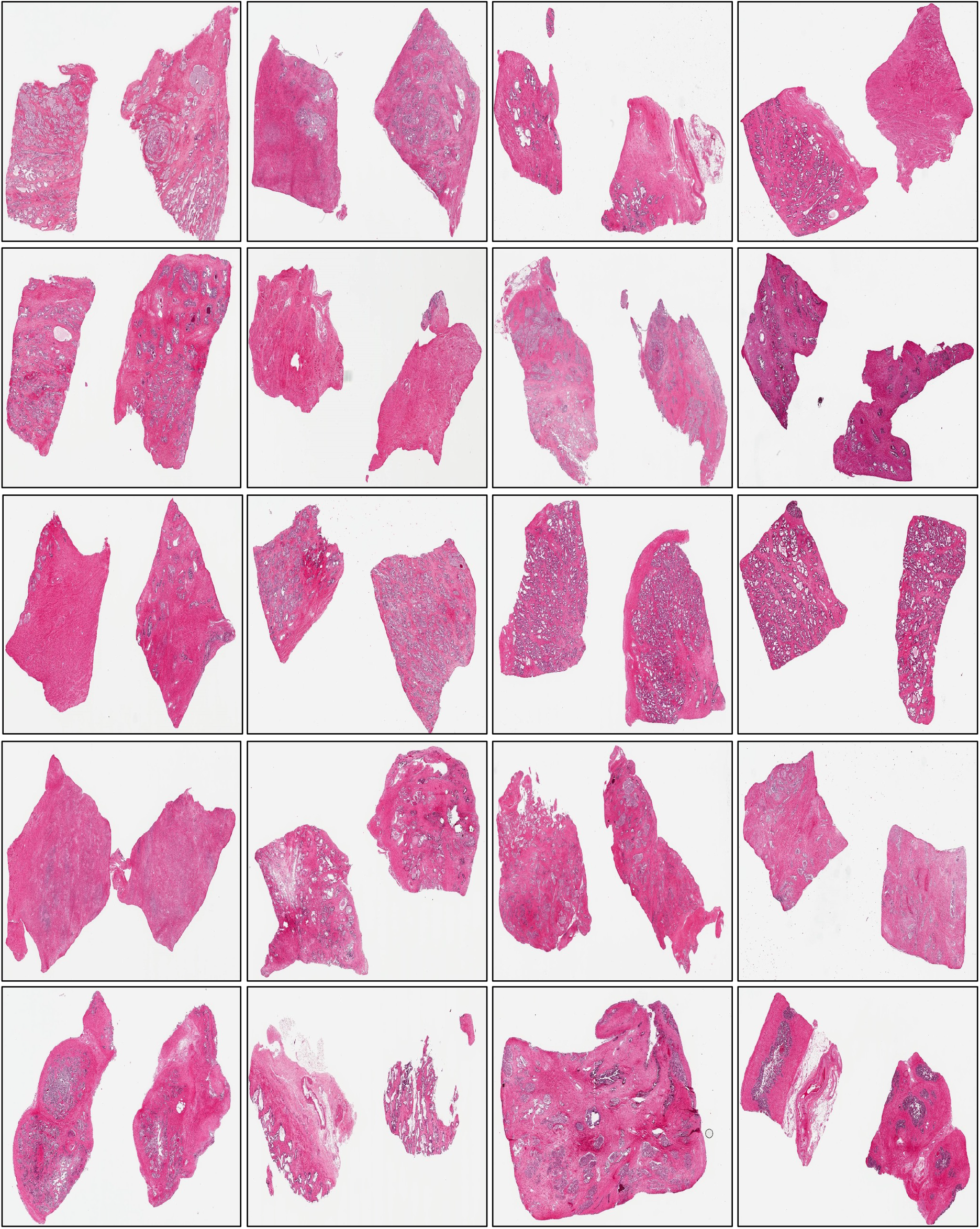
Showing synthetic images of the Prostate.

**Supplementary Figure 6.**
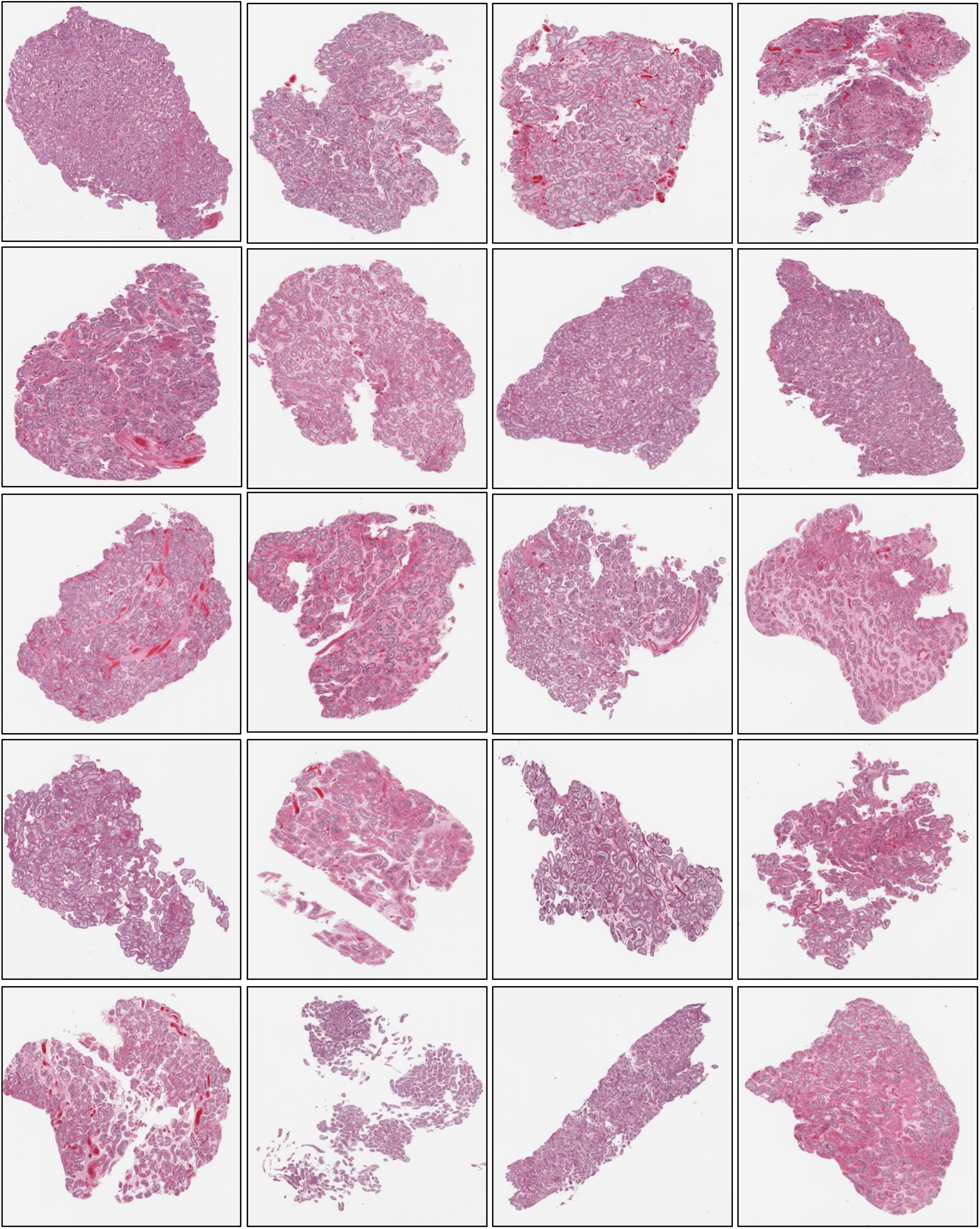
Showing synthetic images of the Testis.

**Supplementary Figure 7.**
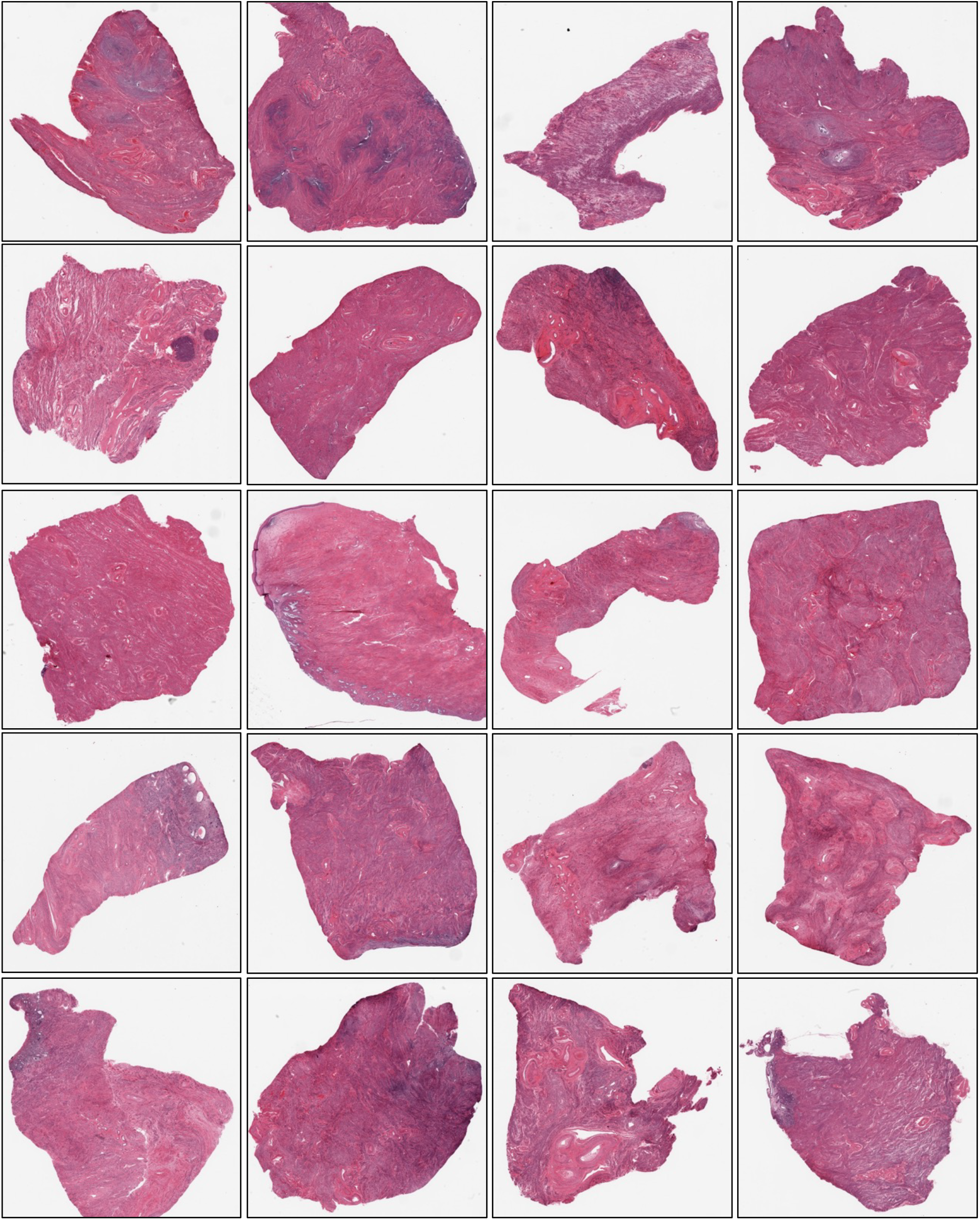
Showing synthetic images of the Uterus.

**Supplementary Figure 8.**
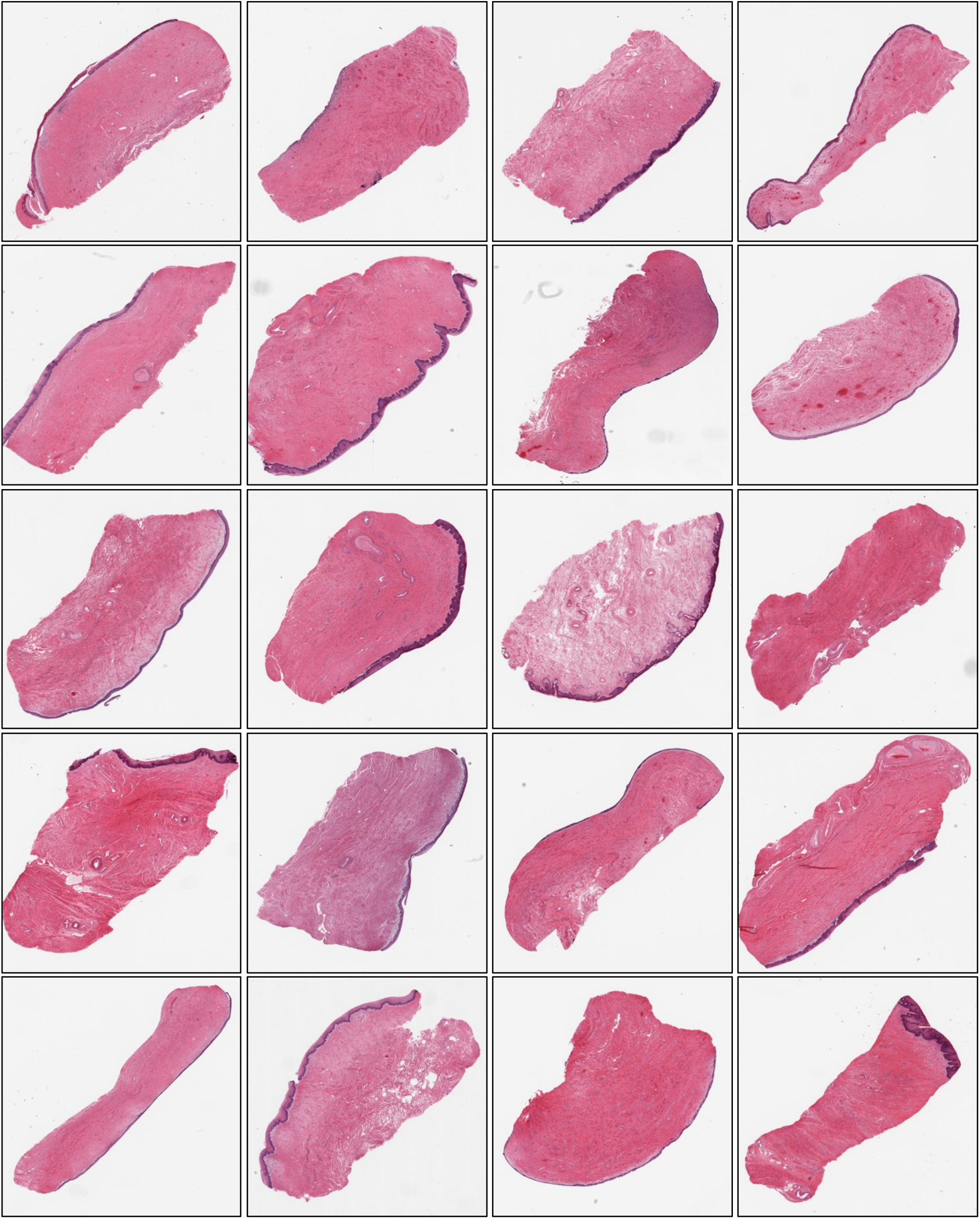
Showing synthetic images of the Vagina.

